# Structure for Energy Cycle: A unique status of Second Law of Thermodynamics for living systems

**DOI:** 10.1101/314567

**Authors:** Shu-Nong Bai, Hao Ge, Hong Qian

## Abstract

Distinguishing things from beings, or matters from lives, is a fundamental question. Extending E. Schrödinger’s *neg-entropy* and I. Prigogine’s *dissipative structure*, we propose a chemical kinetic view that the earliest “live” process is essentially a special interaction between a pair of specific components under a corresponding, particular environmental conditions. The interaction exists as an inter-molecular-force-bond complex (IMFBC) that couples two separate chemical processes: One is the spontaneous formation of an IMFBC driven by the decrease of Gibbs free energy as a dissipative process; while the other is the disassembly of the IMFBC driven thermodynamically by free energy input from the environment. The two processes that are coupled by the IMFBC were originated independently and considered non-living on Earth, but the IMFBC coupling of the two can be considered as the earliest form of metabolism: This forms the first landmark on the path from things to a being. The dynamic formation and dissemblance of the IMFBCs, as composite individuals, follows a principle designated as “… structure for energy for structure for energy…”, the cycle continues, shortly “structure for energy cycle”. With additional features derived from an IMFBC, such as multiple intermediates, autocatalytic ability of one individual upon the formation of another, aqueous medium, and mutual beneficial relationship between formation of polypeptides and nucleic acids, etc., the IMFBC-centered “live” process spontaneously evolved into more complex living organisms with the characteristics one currently knows.

## INTRODUCTION

“To be, or not to be: that is the question”. While Hamlet contemplated life and death in this soliloquy, it is perfectly fitting from the perspective of humanity to question the fundamental distinction, if any, between an animate living being and inanimate non-living matters we call things. The pursuit of the scientific understanding of the origin of life on earth cannot be completely dissociated from the semantics of “what is life”; but as physicist Walker Walker (2017) clearly stated in a highly succinct recent review: “Definitions should emerge from theories, not the converse.” Also very recent in an essay, Steven Rose, an Emeritus Professor of Biology and Neurobiology at the Open University and Gresham College London, summarized (Rose, 2016)that “Modern biology, at its conception in the 17th century, inherited one unshakeable believe, two mysteries and an unfortunate error of timing”. One of the mysteries was over what it is about life that distinguishes it from non-life. Rose believed that this mystery had been solved “by answering that creatures were animate rather than inanimate because there were infused with the breath of life”. He was right to point out that the distinction of life from death is a big mystery in modern biology; but the above answer is only tautological. After all, what is the “breath of life”? Without clarifying this key notion, it is far from conclusive that even the first mystery is solved.

Actually, long before the birth of modern biology and beyond the field proper, efforts had been devoted to draw a line that separates beings from things. The schools of thoughts range from theoretical reasoning, such as Aristotle’s teleology, vitalism proposed by Grancis Glisson, Marcello Malpigi and Caspar Friedrich Wolff (https://en.wikipedia.org/wiki/Vitalism), Oparin’s theory of the origin of life (Oparin, 1953) and its quantitative elaboration proposed by Dyson (1999), Schrödinger’s (1945) celebrated “what is life”, Blum’s effort to correlate the 2nd law of thermodynamic and organic evolution (Blum, 1951), Eigen and Schuster’s (1979) hyper cycles, and Smith and Morowitz’s (2016) “Metabolism First”; to empirical explorations, such as Miller’s synthesis of amino acid (Miller, 1953; Miller and Urey, 1959), Orgel (2004), Wachtershauser (2006), and Copley’s (2015) investigation on prebiotic chemistry.

The various thoughts embody different perspectives. Life can and should be explained from either reductionism or holism. From the former, all matters can be divided into smaller parts. Therefore, living organisms are considered as a combination of atoms and molecules interacting in particular ways. By using the language of chemical reaction kinetics, this approach is successful to describe and predict many biological processes, even complex ones such as embryonic pattern formation (Harrison, 1993). From the holistic standpoint, emergent properties and symmetry breaking were considered to be key necessary but not sufficient ingredients in living systems (Anderson, 1972). However, since nearly all the concepts and methods along this line of inquiry are developed from studying non-living systems, and the purpose of physicists is to discover universal rules to explain behaviors of complex systems that include both living and non-living systems, no line can be drawn to distinguish “beings” from “things” from this “condensed-matter physics” perspective.

Chemists played a major role in understanding the essence of life. From the synthesis of urea by Wöhler, to the discoveries of enzymatic activity and protein structures (Kohler, 1971, 1972 on Buchner’s contribution; Monod et al, 1965; Thomas 2002 on Perutz’s contribution), to elucidation of DNA double helical structure and gene manipulations (Cohen and Chang, 1973; Jackson et al., 1972; Watson and Crick, 1953). All the discoveries repeatedly demonstrated that living processes are essentially chemical reactions. While problems of how these chemical reactions are coordinated and self-reproduced are well addressed (Eigen et al., 1988; Kauffman, 1969; Kauffman, 2011), most of the efforts started from known amino acids and nucleic acids. Although the abstract notion of dissipative structure provided an explanation of one of the essential features of living organisms, the self-organization in open systems (Prigogin.I et al., 1972; Prigogine et al., 1972), it is still unclear how to integrate the chemical reactions demonstrated in test-tubes into a self-sustainable living systems. More importantly, how the events carried out by non-living things evolve in Nature. Here we noted one important ingredient necessary in Darwinian evolution theory: individuals with variations within a population.

In addition to self-organization, self-replication, and dynamics of dissipative chemical reactions, one unique phenomenon for living system is genetic information encoded by a rather permanent, e.g. very stable template called DNA. Some researchers define lives as materialized genetic information (Crick, 1981). This view had dominated modern molecular biology ever since the 1950s. But in recent years, it has gradually met with growing dissenting voices (Noble, 2006). Furthermore, more basic question concerning “what is information” has also been seriously arisen (Qian, 2017).

Beyond the genome, Anfinsen (1973) had proposed that for most single domain proteins, the three-dimensional (3D) folded structure has a minimum free-energy, and such structure is sufficiently determined by the amino acid sequence of a polypeptide. Li et al (1996) showed that a protein’s thermodynamic stability and topological structural fold, called “structural regularities”, could be selected via changes in amino acid sequences in Darwinian evolution. These findings together suggested that the particular sequence of a functional protein may indeed be determined by its biological function subjected to thermodynamic stability of its 3D structures, the fundamental idea of a “folding funnel” (Chan, 1995). From this perspective, thus, DNA sequences just function as records of the biologically active polypeptide sequence with minimum free-energy. But again, this merely provided an alternative explanation to the question of “why something exists” based on Darwinian teleology. It does not give a plausible mechanism for “how” it started (Kirschner and Gerhart, 2005).

Here, we follow the very basic notions of Schrödinger’s neg-entropy (e.g., free energy) and Prigogine’s dissipative structure and move one step further, to explore whether it is possible to identify the first stage, among a series of stages that distinguish beings from things, through an analysis of hypothetic interaction of simplest carbon-based components. The basic new ingredient is to identify a “molecular complex” as an individual ^1^. We believe the notion of individuals is paramount to go beyond the previous physical and physicochemical considerations. It is also clear that with the notion of individuals and population of individuals, biological narratives start. According to the model proposed as follows, what can be called “individual beings” at this very early stage are essentially a special interaction between two specific components under a particular corresponding environmental conditions. Such interaction exists as an inter-molecular-force-bond complex (IMFBC) that couples two independently originated processes. One of the processes is the spontaneous formation of IMFBCs driven by decrease of Gibbs free energy as a dissipative process; while the opposite is the disassembly of the IMFBCs driven by the input of environmental free energy. The dynamic formation and disassembling of IMFBCs that couple the two independently originated processes can be considered as the earliest being with metabolism, follows a principle designated as “structure for energy cycle”. It stands for “…structure for energy for structure for energy…”, the cycle goes on continuously, so is life. Together with the autocatalysis for covalent bonds, derived from the novel structure of IMFBC, and with the aquarium medium, the cycle signifies a natural emergence of the notions of “alive” and “dead” of individuals and the notion of a “continuous living population” in the exactly same context (Huang et al., 2017).

## RESULTS

### The Energy Relationship That Enables the Spontaneous Formation of the IMFBC

Before describing the coupled processes in the structure-for-energy cycle, five elements were considered as necessary conditions: 1) at least two kinds of carbon-based small molecules (generally represented as “components”) used as building blocks for forming the “IMFBCs”; 2) relatively high concentrations of each component in an appropriate niche; 3) formation of the IMFBC from the components; 4) inter-molecular-force(IMF)/interactions for the IMFBCs formation; and 5) input of energy (which can be in the form of chemical free energy) from environment. These first three are conditions concerning “matters”, and the last two are requirements on “thermodynamics”.

With the above mentioned five elements, we hypothesize a scenario in which either the concentrations of components were significantly high, or the energy status of the IMFBCs consisted of the components are significantly lower than that the components exist alone. In such a situation, following the 2nd law of thermodynamics, the self-assembling/forming process from the components A and B to the complex AimB

(IMFBC), in which “im” stands for inter-molecular forces or interactions, would occur spontaneously, as long as the free energy difference

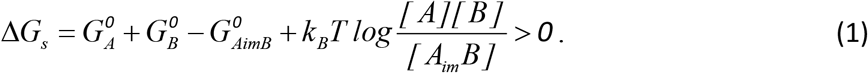

The bracket means concentration or activity, and the *G^0^*s (e.g. *G^0^_A_*, *G^0^_B_*, etc.) are intrinsic free energies.

After a sufficiently long time, in the absence of other processes, the concentration of the complex AimB would stop increasing and the whole system came to an equilibrium state. If this is the case, there would be just a self-assembly process observed in non-living matters such as crystallization. However, if there is certain input of energy from environment to break the inter-molecular interactions, the IMFBCs could be pushed to be disassembled through a distinctly different pathway. During this process, the free energy change should be described by

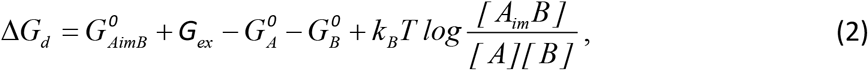

in which G_ex_ is the external free energy input from the environment. One of the easily imagined sources of G_ex_ is solar energy or simply heat.

Overall, the total free energy driving the process composed by the forming and dissolving steps was

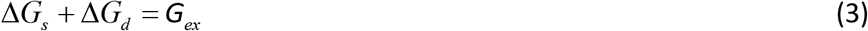

Therefore, in the presence of external free energy *G_ex_* > *0*, the net fluxes of the forming and disassembling steps could be both positive, constitute a cyclical process. The total concentrations of the components as well as the IMFBCs would not change with time at steady state, but since the forming and disassembling went through different pathways, it was a non-equilibrium steady state with non-vanishing metabolism rate (Qian, 2006). This might be the feature so unique that enables us to refer such a cyclic process the first live system, or “being”.

Such a cyclical process could be represented as diagram such as Figure 1.

**Figure 1.**
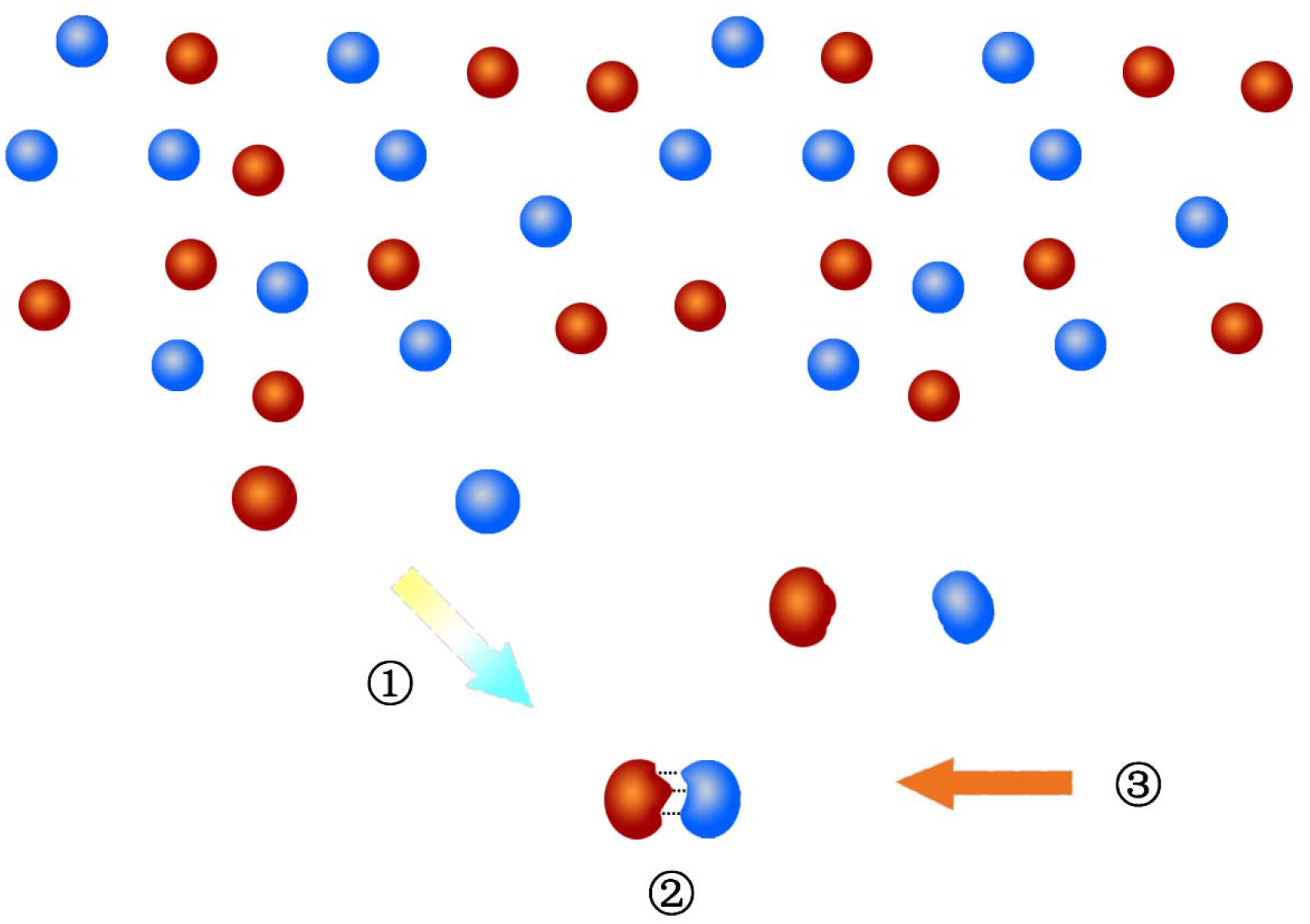
The diagram of a non-reversible cycle of two independent processes coupled by IMFBC, designated as the first live system. Red and blue balls represent particular carbon-based components with different features. Colored arrow (circled 1) represents a process of which a spontaneous formation of a IMFBC (circled 2) driven by the reduced free energy. The light brown arrow (circled 3) represents an input of energy from corresponding environment, which breaks the inter-molecular-force and dissolves the IMFBC.

The driving forces of the two processes are different. The driving force for the first mainly comes from the difference of the structures and concentrations between the freely existing components and the IMFBCs, in which, the former has higher chemical potential than the latter and thus the “formation process” (synthesis) can spontaneously occur. In contrast, the driving force for the second process (degradation) mainly come from the input of external energy, which break the weak inter-molecular forces used for the formation of the IMFBCs and release the components.

Due to the fact that the free energy input from the environment cannot be extremely high and a reasonable living entities needs to be flexible within reasonable time scales, the free energies of both transition states along the formation and disassembling steps cannot be extremely high, especially at the ancient times without proper enzymes as catalysts.

Because the uniqueness of the processes comes from the formation of the IMFBCs with particular structure which is bonded by inter-molecular interaction, and the IMFBCs formed through the reduction of free energy and disassembled by the input of external free energy, the principle/rule/law underlies the processes described above was designated as “structure for energy cycle”.

### The Energy Relationship That Enables the Formation of Covalent Bonds

However, although the cyclic process described by the formula 3 defined “being” and therefore drew the first line between living and non-living processes, it simply remains as a rather uninteresting circulation to a biologist, and one of common chemical reactions occasionally occurs that is never being considered relevant to living to a chemist. Without covalent bonds, there would be no real bio-molecules and there would be no sufficiently stable “biochemistry” that is evolvable. The non-covalent scenario is highly environmental sensitive. Covalent bonds make the labile complexes much more stable and robust. But how are the covalent bonds introduced into this beginning origin of a living system?

Thermodynamically, the process of the formation of covalent-bonded complexes can be described as follows:

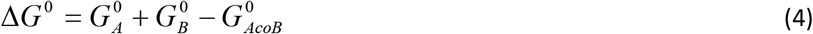

in which AcoB refers to the complexes formed by covalent bonds.

The energy barriers of covalent bonds are much higher than that of inter-molecular non-covalent bonds, so is the stability (Figure 2). Therefore, the formation of covalent bond is anticipated to consume much more energy than the formation of intermolecular interaction. The Miller-Urey experiment on synthesis of amino acids from inorganic precursors under conditions of higher temperature together with electric sparks is a key experimental fact that motivated our hypothesis (Miller and Urey, 1959; Saita and Saijia, 2014).

**Figure 2.**
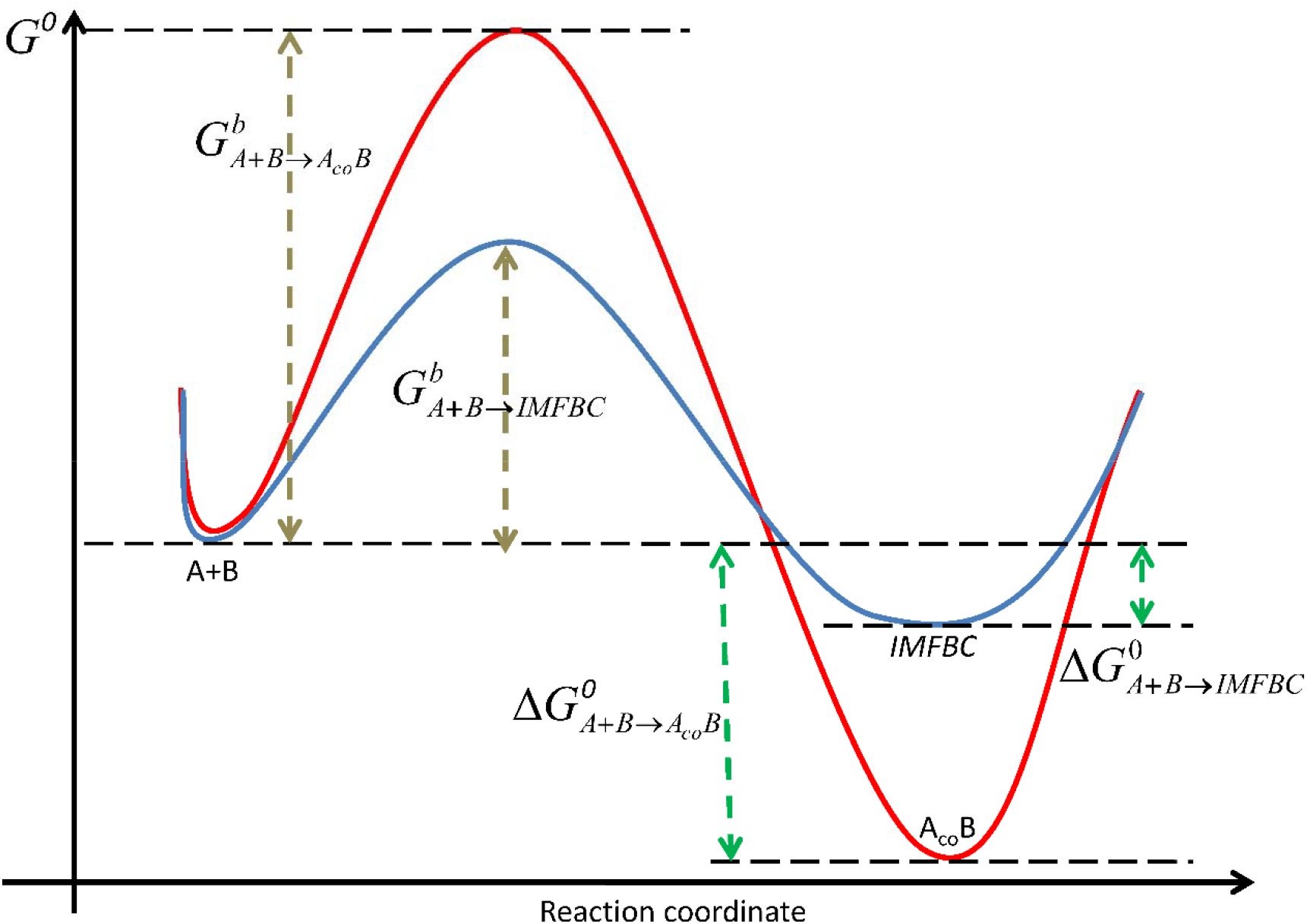
Free energy surfaces for the synthesis of IMFBC (blue) and A_co_B (red) The energy barrier for the synthesis of IMFBC (GbA+B->IMFBC) is much lower than that for the synthesis of A_co_B (GbA+B->AcoB).

It is known that current living systems on the Earth consist of carbon-based components. It is believed that the tetravalence at the second electron shell enables carbon to form large complex molecules. If the above proposed “live” process indeed occurs, it is obvious that the carbon-based components that consist of the IMFBC (AimB) should have extra space available for additional bonds to form. If the IMFBCs happened to have a particular surface with autocatalysis features to reduce energy barrier for covalent bond formation, or happen to be mixed with some catalytic surfaces such as FeS2 as suggested by Wachtershauser (1988), these IMFBCs can function as a reaction active center for spontaneous covalent bond formation with other carbon bone components at the presence of the inter-molecular interaction remains. Figure 3 describes the hypothetic process for spontaneous covalent bond formation upon the preexisting IMFBCs (AimBs).

**Figure 3:**
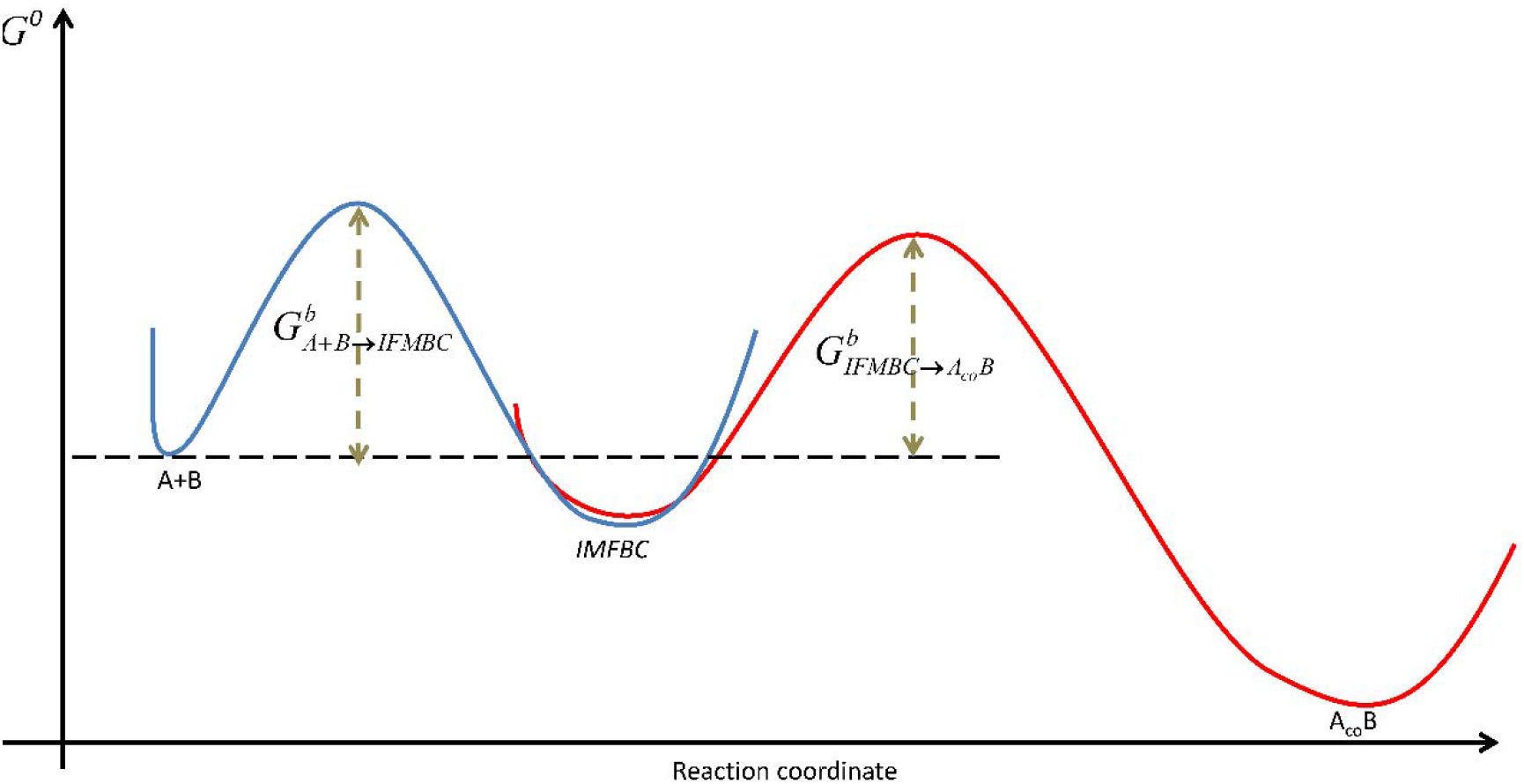
Surface of IMFBC may reduce energy barrier to form covalent bonds. Under the hypothetic surface autocatalysis of preexisting IMFBC, the energy barrier for IMFBC converting to A_co_B (red) is much lower than that for A+B to directly form covalent bonds inside A_co_B (red in Figure 2).

Through the process described by Figure 3, in the same system described by Figure 1, if the system is open not only with input energy to disassemble the spontaneously formed IMFBCs, but also input materials to keep the population of the components, the preexisting IMFBCs (AimBs) will increase their complexity. The hypothetic surface autocatalysis of the IMFBCs (AimBs) explains not only the spontaneous formation of covalent bond with the preexisting IMFBCs (AimBs) as reaction center, but also the increase of complexity of bio-molecules. This feature breaks the boring circulation described in formula 3 and Figure 1 and essentially initiates a path of evolution.

### Additional Components Required for the Formation of a Living System

Formula 3 and Figure 1 described a “live” process that drew the first line to distinguish living and non-living system. The structure-for-energy cycle can be therefore considered as the first hallmark of living system. Figure 3 explained how a covalent bond spontaneously formed upon the preexisting IMFBCs (AimBs) through autocatalysis and how “evolution”, e.g. increase of complexity, originated and become the second hallmark of living system. However, it is common sense that living systems are far more complicated than the two hallmarks. For example, water is not required for both formations of IMFBCs and covalent bonds. How to explain indispensability of water for living system? More challenging issues are the origin of genetic code and emergence of networks consisting of more sophisticate metabolic and signaling pathways.

At what point water becomes indispensable to a living system? If the spontaneous covalent bond formation indeed occurred because the autocatalytic feature of the surface of IMFBCs, there would be two problems to be solved. One is how to prevent aggregation of spontaneously formed molecules, and the second is how to buffer the external energy input to maintain a proper niche for the live process to evolve. Probably, water was selected as an ideal medium to solve both problems because its unique features such as being a powerful solvent and having big specific heat capacity. Together with other functions, from solving the above two problems on, may water become an indispensable component in the splendid evolutionary journey of living system.

How the relationship between polypeptides and nucleic acids was established? It is a long lasting puzzle. Various hypotheses have been proposed to explain how the genetic code origin (Hopfield, 1978 and reference therein). It is reasonable to propose that in the “primordial soup”, the IMFBCs with different carbon-based components are formed randomly. While the energy status considered, it is again reasonable to hypothesize that amino acids can bond together to form polypeptide, and nucleotides bond together to form nucleic acid from the perspective of energy favorability. However, Drygin (1998) had reported that molecules consisting of both nucleotide and amino acids can be naturally formed. Copley et al. (2005) had reported the synthesis of amino acids with dinucleotides complex covalently. These phenomena suggested interactions between the formation of amino acids/peptides and nucleic acids in the “primordial soup”. Such interactions may imply inter-catalysis of the two types of macromolecules. Taken the discovery of Li et al (1996) in consideration, it is possible that the sequences of polypeptides were originally determined by the energy favorability. Such energy favorable sequences were recorded by nucleic acid sequences through the interaction between the polypeptides and nucleic acids formations, probably the catalytic effects of the forming polypeptides upon the nucleic acid formation. In turn, decoding of the nucleic acid sequence to polypeptide formation will significantly increase the efficiency to reproduce the energy favorable polypeptide sequences. Among such a complicated interaction, tRNA may play a critical role as suggested by Hopfield (1978). Along with the numerous rounds of selection, although with inevitable increase of complication, the information of energy status of polypeptide sequences was recorded into base pair sequences of nucleic acids, and nucleic acid, especially in the form of most stable molecule DNA, become a storage center for the energy favorable polypeptide sequences of entire living systems, and further become a hub of the complicated macromolecule networks. as described by the “central dogma” (Crick, 1958; Crick, 1970).

Thus far, five processes were described, including a) “live” process centered with the IMFBCs; b) autocatalytic surface of IMFBCs for covalent bond formation; c) integration of water into the living system as an indispensable component; d) energy favorability determined polypeptide sequence; and e) the mutual benefit of polypeptide and nucleic acid interaction. If those five processes exist in reality, it seems ready to spontaneously form networks consisting of sophisticate metabolic and signaling pathways for emergence of the living system with characteristics we currently knew. One additional component required for the network formation would be the catalytic feature derived from some polypeptides, i.e. enzymatic activity, that facilitates the formation of other macromolecules, under assistance of high energy molecule, such as ATP and GTP etc.

Another insight from the current theory is the realization that while the form of a live persists, the living matter is only transient: This is because in a nonequilibrium steady state (NESS), while the population of the IMFBC continues, the individuals within the population only exists for a period of time that is related to the rate parameters in the disassembly process. This *stable form with unstable matters* is a fundamental character of mesoscopic reversible chemistry as well as living organisms. The former is sustained by detailed balance while the latter by synthesis and degradation.

## DISCUSSION

While it is widely considered that life started from a cell, some fundamental characteristics of living systems also appear in cell-free circumstance. Therefore, it is not unreasonable to draw an earlier line between beings and things to pre-cellular stages. Differing from the mainstream efforts to explore origin of life through analyzing the formation of currently known molecular building blocks, e.g. material basis such as amino acids and nucleotides, here we continue the approach pioneered by Schrödinger and Prigogine, via physicochemistry, by emphasizing the role of inter-molecular force (IMF) in the emergence of living matters: they cannot be understood as equilibrium matter alone but as a dynamically balanced formation and dissembling cycle. The rationale for this approach is simple. Behind all the characteristics unique to currently known living systems are specific IMFs. These characteristics include base pairing in DNA double helix, via hydrogen bonds and base-stacking; active sites of enzymes and overall functional structures of proteins maintained by hydrogen bonding and van der Waals interactions; and allosteric effects resulting from the combination of IMFs which sometimes enhanced by the involvement of metal ions. The IMFs are traditionally considered as properties of macromolecules. But according to our analysis, as described in formula 3 and Figure 1, IMFs could play a key role in forming an IMFBC, and endows the IMFBC with a central role in coupling the two independently originated processes, which enables macromolecules to emerge spontaneously.

Briefly, in a particular moderate environment with which the five conditions, i.e. “carbon-based components”, concentration, IMFBC, IMF and openness to environment, were simultaneously met, two spontaneously occurred processes can be coupled as ONE cyclical process follow a rule called “structure for energy”, as a unique status of the Second Law of Thermodynamics. Such a cyclical process can be defined as the first “being” or “living matter”, thus drew the line to distinct “being” and “thing”. It has been argued that there are many chemical reactions possess characteristics similar to the IMFBC-centered cycle and therefore, the IMFBC-centered cycle should not be considered as the first “being” or “living matter”. Indeed, only based on a cyclical property, the chemical reactions are not qualified to be considered as a “living organism”. However, the uniqueness for the IMFBC-centered cycle we hypothesized is that one should not consider a material entity alone as alive, but rather as a start, such an entity involved in a dynamic cyclic process that is driven by external energy. We believe this notion is novel; it is certain derived from the particular feature of the carbon-based components, the IMFBCs are spontaneously with autocatalytic activities of the surface. Only the IMFBCs with such characteristic, the covalent bonds can be spontaneously formed, which leads to increase of complex that is the essential feature of evolution. Without the IMFBC, there might be cyclical chemical reactions, but would be no spontaneous increase of complex accompany with the cyclical process. That is why we referred the IMFBC-centered “live” process as the first line of “being” from “thing”. With IMFBC as the first necessary condition, the autocatalysis of the IMFBC as the second and the water medium as the third, a robust living system emerges. Further, with mutual beneficial relationship between formations of polypeptides and nucleic acids, the IMFBC-centered “live” process spontaneously evolved into more complex living systems with characteristics we currently know. Since the key steps described in the early evolution of living process were derived on particular structures as well as specific processes, such a view on the origin of life can be therefore designated as “Structure and Dynamics Cycle First”. The concept of information is simply a type of probabilistic or biological interpretation of the cyclic process (Qian, 2017).

Among the efforts to explore the fundamental rules governing living systems, one approach is to treat “being” as a system and tried to discover the rules from mathematic description of the overall behavior of the system. Qian et al. (2016) have proposed that living systems, not very different from P. W. Anderson’s hierarchical structure of nature (Anderson, 1972), necessarily exhibit different features at different scales. The IMFBC-centered live process is consistent with a stochastic, microscopic molecular world and provides a plausible molecular mechanism. From this perspective, the reductionism and holism are not mutually exclusive. If one finds a proper point to start with, both perspectives will be mutually beneficial. Note our theory as all thermodynamic assertions can only tell what is impossible and thus what a real possibility has to respect. It does not suggest a more specific mechanism. At this point, however, our theory makes a logic contact with the epic scenario presented by Smith and Morowitz (2016): Based on a sophisticated scientific analysis of geochemical status, and the chemical synthesis feasibility, they arrived at the conclusion that carbon metabolic cycle is the origin of emergent life. In the language of our theory, their entire rTCA cycle can be considered as a single IMFBC, with sufficient realism. We note that the synthesis and degradation of an IMFBC, as two distinct chemical processes, have to pass different intermediate states: There must be a multi-stage cycle in chemical details. This inference is consistent with Morowitz’s (1968) cycling theorem, which in turn is in complete agreement with the modern theory of NESS (Zhang et al., 2012)

The theory of NESS (Zhang et al., 2012) also unifies our view of evolution in a stationary chemical environment with given chemical potential difference(s) and Smith-Morowitz’s view of entire geophysical chemistry with a continuous energy flow. In the latter perspective, they argued that the emergence of rTCA type of biochemistry is a necessity driven by the flow of electrons from higher to lower potentials, coupling geochemistry with biosphere on earth. Finally, but not the least, we point out the “phase transition paradigm” forcefully articulated by Smith and Morowitz is hidden in our theory: The chemical association, i.e., recognizing AB complex as an individual entity in a sea of A and B molecules in a solution, is a phase transition in atomic physics, but a matter of fact to high-school chemistry (Fisher and Zuckerman, 1998). Indeed, as poignantly said in Smith and Morowitz (2016): “Chemistry unifies the extraordinary diversity of living order to a degree that no other starting point can”.

Let us recap what is our novel proposition, if any. As already clearly recognized by Schrödinger in 1945, and forcefully argued by Blum in 1951, the openness of a chemical system is one of the fundamental aspects of lives constrained by physical laws. More importantly, an open chemical system has to be situated in a complex environment that has at least *two* different thermodynamic potentials (Blum, 1951), a notion further advanced in the theory of dissipative structures. We argue that the next logical conclusion is that the live as a phenomenon can no longer be fully understood as an entity along, but an individual within a cyclic chemical reaction system that at least has two distinct processes. The individual starts as simple as an IMFBC (inter-molecular-force-bond complex) and evolves into as complex as a unicellular organism; the corresponding processes are as simple as structural formation and dissemblance, and become more complex hype-cycles with both auto-catalysis and self-replications (Eigen and Schuster, 1979). One of the emergent aspects of such a cyclic open chemical reaction system is the continuous interaction between the individual in terms of its own structure and the energy that is present in the environment: This, we argue, is the fundamental aspect of *metabolism*. It is interesting to note that the very term “metabolism” in non-English languages, e.g., in German Stoffwechsel (Dyson, 1999) and in Chinese Xīnchéndàixiè 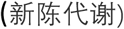, simply means “structural renewal through chemical processes” without any connotation in connection to replications and genetics.

A second, subtler conclusion can also be obtained logically, if one recognizes two very different “chemical view” vs. “mechanical view” of the Natural World (Qian et al., 2016; Fisher and Zuckerman, 1998). In the latter, an individual is a point mass which is featureless with only mass, position, and velocity as its traits. However, in a chemical view of the Natural Worlds, individuals are atoms and molecules with infinitely complex constituents, if only one probes deeper. Under the chemical perspective, two particles with affinity can form a complex is considered as self-evident, while in mechanics, it requires a theory to elucidate the emergence of a new “quasi-particle” as a consequence of a phase-transition like process (Walker, 2017; Smith and Morowitz, 2016).

## ACKNOWLEDGEMENTS

We thank Professor Ping Chen of Fudan University for his encouragement to write down an idea into this paper; Professors Xiao-Dong Su, Yi-Qin Gao, Zhi-Rong Liu, and Xin-Sheng Zhao of Peking University for their inspiring discussions on various related issues. We also thank G. Allen (Washington Univ. St. Louis), R. H. Austin (Princeton), Y. Chen (ENS Paris), H. Li (UCSF), Q. Ouyang (PKU), D. E. Smith (SFI), Y. Tu (IBM), R. A. van Santen (TU Eindhoven), X. Yu (Inst. Neurosci., CAS), C. Zhang (Inst. Biophys., CAS for their reading of the manuscript and helpful comments. This project was supported by MST 2003CB715906 to SB and NSFC 11021463 to OYQ.

## Footnotes

1 The term “individual” originally meant to be “indivisible”. This is not what we imply when we employed this term here. Rather, we use the term “individual” as opposite to “population” as two essential terms in evolutionary biology. It is important to recognize that a “living individual” actually is a “dynamic form” within which the materials are continuously changing anew. In the current context, the atoms in an IMFBC complex are not the same due to continuous assembly and disassembly.

